# The Choice of Reference Frame Alters Interpretations of Non-Linear Gait

**DOI:** 10.1101/2022.11.01.514761

**Authors:** Tyler K. Ho, Nicholas Kreter, Cameron B. Jensen, Peter C. Fino

**Affiliations:** University of Utah, Department of Health and Kinesiology, Salt Lake City, UT, USA

**Keywords:** Turning, locomotion, margin of stability, gait stability, coordinate system, motion capture

## Abstract

**Introduction:** Humans regularly follow non-linear trajectories, such as turning, during everyday ambulation. However, globally-defined and locally-defined reference frames fall out of alignment during non-linear locomotion, which complicates spatiotemporal and biomechanical analyses of gait. Thus, the choice of the locally-defined reference frame is an important methodological consideration. This study investigated how different definitions of reference frame change the results and interpretations of common gait measures.

**Methods:** Nine healthy adults completed two walking trials around a circular track. Kinematic data were collected via motion capture and used to calculate step length, step width, anteroposterior margin of stability, and mediolateral margin of stability using three different locally-defined reference frames: walkway-fixed, body-fixed, and trajectory-fixed. Linear-mixed effects models compared the effect of reference frame on each gait measure, and the effect of reference frame on conclusions about a known effect of turning gait – asymmetrical stepping patterns.

**Results:** All four gait measures differed significantly across the three reference frames. A significant interaction of reference frame and step type (i.e. inside vs outside step) on all four gait measures (*p* < 0.001) indicated conclusions about asymmetry differed based on the choice of reference frame.

**Conclusion:** The choice of reference frame will change the calculated gait measures and may alter the conclusions of studies investigating non-linear gait. Care should be taken when comparing studies that used different reference frames, as results cannot be easily harmonized. Future studies of non-linear gait need to justify and detail their choice of reference frame.

## 1. INTRODUCTION

Defining the reference frame of a locomotor environment is a critical step in reporting consistent and comparable measures of gait, with internationally accepted standards defined during straight gait (Wu and Cavanagh, 1995). However, humans routinely walk in non-linear trajectories where there is no clear, pre-defined direction of travel (Glaister et al., 2007a). Compared to straight-line walking, spatiotemporal and biomechanical analysis of this non-linear gait is more difficult to analyze because the globally-defined and locally-defined reference frames fall out of alignment (Huxham et al., 2006; Kainz et al., 2016). In these non-linear trajectories, measures from globally- and locally-defined reference frames can vary dramatically, yielding inconsistent interpretations and conclusions based on the choice of coordinate reference frame (Schache et al., 2008). Several different locally-defined references frames have been used to study non-linear gait such as turning. But, there are no guidelines on how to choose a locally-defined frame (Glaister et al., 2007b; Wu et al., 2002; Wu and Cavanagh, 1995), or whether results from different locally-defined reference frames can be combined, despite non-linear locomotion being increasingly studied.

To help provide guidance when choosing a locally-defined reference frame for non-linear / turning gait, the aim of this study was to investigate how the choice of reference frame can influence spatial measures of gait and affect conclusions about important effects of interest during a non-linear gait (i.e., turning) task. To do so, we used positional marker data to calculate step length, step width, anteroposterior (AP) margin of stability (MoS), and mediolateral (ML) MoS in three different reference frames: walkway-fixed, body-fixed, and trajectory-fixed. Asymmetrical gait patterns have been well-described during turning (Fino et al., 2015; Orendurff et al., 2006; Strike and Taylor, 2009), so we examined how the different reference frames affected conclusions about asymmetries during turning. We hypothesized that the choice of reference frame would change the resulting gait measures discussed above, and that such differences would affect the conclusions about asymmetrical stepping patterns during turning gait. Finally, we provide recommendations for interpreting results across past studies and selecting an appropriate reference frame for future non-linear gait research.

## 2. MATERIALS and METHODS

### 2.1 Participants

Nine healthy adults [5 female, 4 male; mean (SD) age: 22.9 (3.1) years; height: 172.6 (10.7) cm; mass: 64.7 (9.7) kg] were recruited from the local community and provided informed written consent for participation in this IRB-approved study. Exclusion criteria included: (1) any neurological condition that could affect balance, (2) unresolved musculoskeletal injury to the lower extremities, (3) any reconstructive surgery to the lower limbs, or (4) an inability to withhold prescribed or recreational pharmacotherapy, that could affect balance or psychological state, for 24 hours surrounding data collection.

### 2.2 Procedure

Participants completed walking trials around a 0.4m wide circular track (inner radius: 1.2m, outer radius: 1.6m) constructed of 2.5cm thick plywood. The raised track ensured no participant deviated outside of the inner or outer radius. As part of a larger protocol, the participants completed one walking trial in the clockwise direction and one in the counterclockwise direction, for one minute each. Participants’ self-selected speed while walking around the track was measured prior to the first trial, and they were paced with a metronome delivered at each quarter-lap to maintain a consistent walking speed throughout each trial. Retroreflective markers were placed bilaterally on the posterior superior iliac spine (PSIS), anterior superior iliac spine (ASIS), iliac crest, heel, second metatarsophalangeal (MTP) joint, and fifth MTP joint. Two extra markers were placed on the feet [first tarsometatarsal joint and first MTP joint] of two participants to reduce processing time. Kinematic data were collected via motion capture (Vicon Nexus ver. 2.12) at a rate of 200 Hz.

### 2.3 Data Processing

Custom MATLAB codes were used for data processing and statistical analysis (ver. R2020b, The Mathworks Inc., Natick MA, USA). Raw positional data from all markers were filtered using a 4^th^-order phaseless Butterworth filter, with a low-pass cut-off frequency of 6 Hz. The position of the body center of mass (CoM) and feet in a globally-fixed reference frame were estimated from the six hip markers (Alcantara et al., 2021; Havens et al., 2018) and the heel, second MTP, and fifth MTP markers, respectively. The velocity of the CoM was calculated using the central difference formula. Temporal gait events of heel contact and toe-off were identified according to the methods by Zeni et al. (Zeni et al., 2008), modified for turning gait (Ulrich et al., 2019). Steps were categorized into either inside or outside steps, based on which direction they moved around the circle (e.g., clockwise: right is inside; counterclockwise: left is inside).

Each position and velocity vector was rotated from the globally-fixed reference frame to three different reference frames: a walkway-fixed frame (Wu and Cavanagh, 1995), a body-fixed frame (Schache et al., 2008; Wu et al., 2002), and a trajectory-fixed frame (Kainz et al., 2016) [Figure 1]. The walkway-fixed frame was defined using the radial vector from the center of the circular walkway to the position of the CoM (+*Z*_*W*_), vertical (+*Y*_*W*_), and the right-handed orthogonal cross-product of the two (+*X*_*W*_), with +*X*_*W*_ pointed in the direction of travel. The body-fixed frame was defined by the vector from the average position of the PSIS markers to the average position of the ASIS markers (+*X*_*B*_), vertical (+*Y*_*B*_), and the cross-product of the two (+*Z*_*B*_). The trajectory-fixed frame was defined by the vector of the instantaneous velocity of the CoM in the transverse plane (+*X*_*T*_), vertical (+*Y*_*T*_), and the cross-product of the two (+*Z*_*T*_).

**Figure 1:**
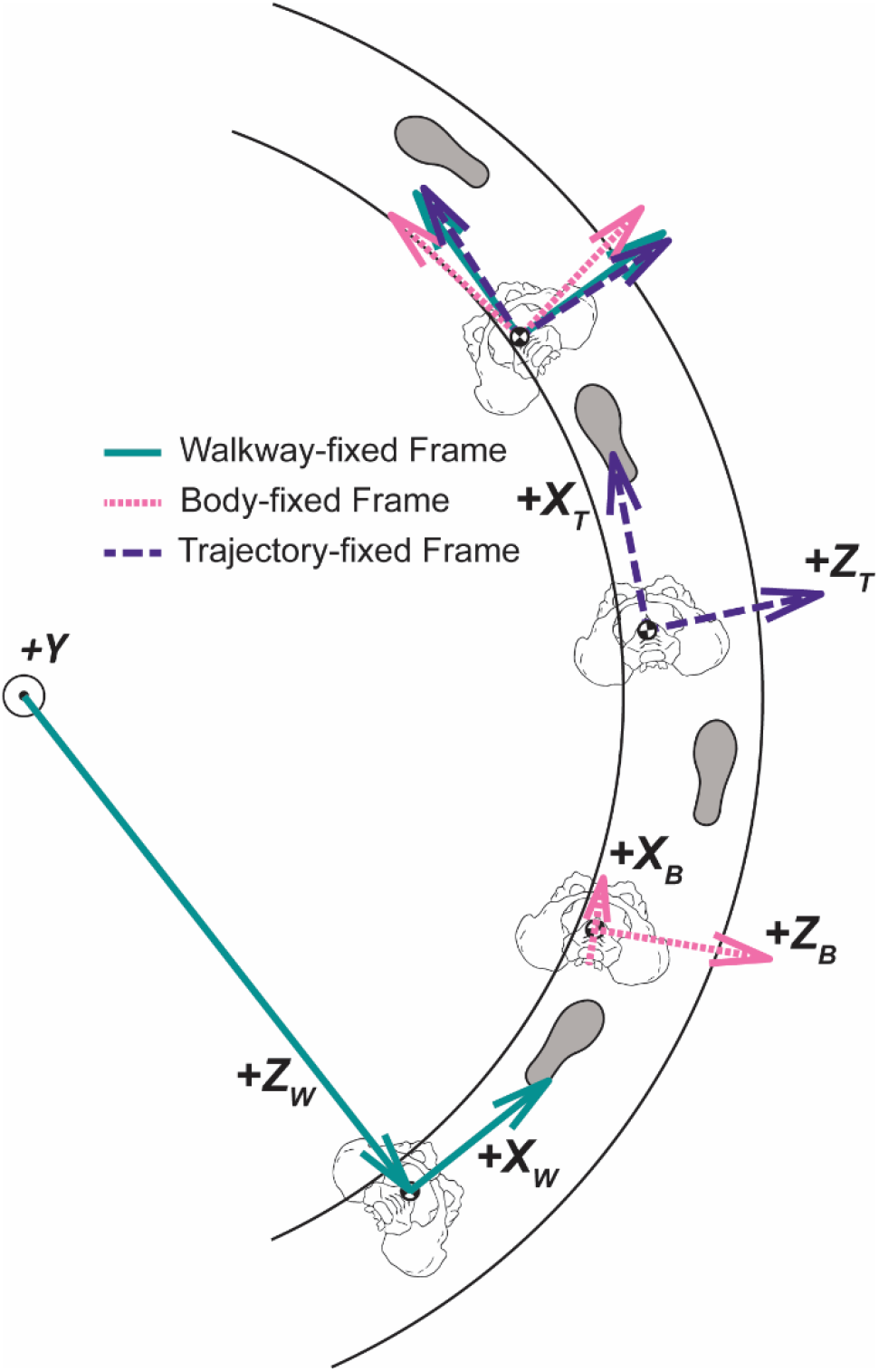
Visualization of the walkway-fixed (green, solid), body-fixed (pink, dotted), and trajectory-fixed (purple, dashed) reference frames. The ML aspect of the walkway-fixed frame (*+Z*_*W*_) was defined by the radial vector from the origin of the circular track to the body CoM. The AP aspect of the walkway-fixed frame (*+X*_*W*_) was defined by the orthogonal cross-product of *+Z*_*W*_ and a vertical vector (*+Y*). The AP aspect of the body-fixed frame (*+X*_*B*_) was defined by the vector from the average position of the PSIS to the average position of the ASIS. The ML aspect of the body-fixed frame (*+Z*_*B*_) was defined by the orthogonal cross-product of *+Z*_*B*_ and *+Y*. The AP aspect of the trajectory-fixed frame was defined by the vector of the instantaneous velocity of the CoM (*+X*_*T*_). The ML aspect of the trajectory-fixed frame was defined by the orthogonal cross-product of *+X*_*T*_ and *+Y*. Each reference frame was calculated at initial heel strike for measuring step length and step width, and at contralateral toe-off for measuring AP and ML MoS.

Step length and step width were calculated as the *X* and *Z* components, respectively, of the vector connecting the heel markers of successive heel strikes (e.g., right to left) within each reference frame, which were extracted at the instant of the initial heel strike. The positive directions for each step pointed in the local *+X* direction and along the medial direction of the local *Z* axis from the initial heel strike (i.e., a negative step width indicates a crossover step). The AP and ML MoS were calculated as the length of the vector from the extrapolated CoM (xCoM) – a measure of the position and velocity of the CoM – to the most anterior and lateral borders of the base of support (BoS), respectively, at the time of contralateral toe-off (Hof, 2008; Hof et al., 2005). The most anterior and lateral boundaries of the BoS were calculated by defining a piece-wise linear boundary from the second MTP marker to the fifth MTP marker and from the fifth MTP marker to the calcaneus marker (2MTP-5MTP-Calcaneus). For each reference frame, separate AP and ML BoS boundaries were defined as the points along that piecewise 2MTP-5MTP-Calcaneus line that were furthest from the xCoM in the local AP and ML directions, respectively. The positive directions for MoS pointed in the local –*X* direction (AP) and along the medial direction of the local *Z* axis from the borders of the BoS (e.g., a negative AP MoS indicates the xCoM is located in front of the anterior border of the BoS, and a positive ML MoS indicates the xCoM is located medially to the lateral border of the BoS).

### 2.4 Statistical Analysis

To test the effects of reference frame on stability measures within subjects, linear mixed models were fit for each outcome measure with fixed effects of reference frame, stepping limb (inside vs outside), and their interaction. Models were also adjusted for the covariate of walking direction (clockwise vs. counter-clockwise). Each model included random intercepts by subject to adjust for within-subject correlation. Overall effects of reference frame and the interaction with stepping limb were examined using a type 3 test for fixed effects (i.e., F-test). Post-hoc contrasts followed the F-tests and tested for pairwise differences between reference frames. For each model, significance was assessed at an alpha of 0.05, and a Benjamini-Hochberg correction (Benjamini and Hochberg, 1995) was used to account for multiple comparisons.

## 3. RESULTS

Step length (*dF* = 4838, *F* = 538.7, corrected *p* < 0.001), step width (*F* = 556.9, corrected *p* < 0.001), AP MoS (*F* = 29.8, corrected *p* < 0.001), and ML MoS (*F* = 109.6, corrected *p* < 0.001) all differed across reference frames [Figure 2A-D]. Altering reference frame also changed the observed asymmetry of stepping limb (i.e., inside vs outside) on all four measures, as indicated by the interaction between reference frame and stepping limb (*F*_*SL*_ = 462.8, corrected *p*_*SL*_ < 0.001; *F*_*SW*_ = 103.8, corrected *p*_*SW*_ < 0.001; *F*_*AP*MoS_ = 119.0, corrected *p*_*APMoS*_ < 0.001; *F*_*MLMoS*_ = 8.0, corrected *p*_*MLMoS*_ < 0.001) [Figure 2A-D]. Pair-wise contrasts failed to detect a significant interaction of reference frame and stepping limb on step width when comparing the walkway-fixed frame to the trajectory-fixed frame (corrected *p* = 0.184) [Figure 2B; Table1]. Pair-wise contrasts also failed to detect a significant interaction of reference frame and stepping limb on ML MoS when comparing the walkway-fixed frame only to the body-fixed frame (corrected *p* = 0.063) [Figure 2D; Table 1]. All other pair-wise comparisons were significantly different (see Table 1).

**Table 1:** Coefficients and results from the linear mixed models for step length, step width, AP MoS, and ML MoS. Fixed effects included reference frame, stepping limb, walking direction, and the reference frame*stepping limb interaction. The walkway-fixed reference frame and inside stepping limb were the reference condition for each model. Each gait measure differed both by reference frame and stepping limb. The interaction of reference frame*stepping limb on step length and AP MoS was significant across all reference frames. However, the interaction of reference frame*stepping limb did not alter interpretations of step width when comparing the walkway-fixed frame to the trajectory-fixed frame, nor did it alter interpretations of ML MoS when comparing the walkway-fixed frame to the body-fixed frame.

|  | **Name (dF = 4838)** | **Estimate (mm)** | **SE (mm)** | **pValue** | **95% CI (mm)** |
| --- | --- | --- | --- | --- | --- |
| **Step Length** | Intercept (Walkway-fixed Frame, Inside Step) | 565.0 | 9.6 | < 0.001 | [546.3, 583.8] |
|  | Body-fixed Frame | 30.5 | 1.6 | < 0.001 | [27.3, 33.7] |
|  | Trajectory-fixed Frame | -22.9 | 1.6 | < 0.001 | [-26.1, -19.7] |
|  | Stepping Limb | 45.6 | 1.6 | < 0.001 | [42.4, 48.8] |
|  | Walking Direction | -0.7 | 0.9 | 0.487 | [-2.5, 1.2] |
|  | Body-fixed Frame*Stepping Limb | -35.7 | 2.3 | < 0.001 | [-40.2, -31.1] |
|  | Trajectory-fixed Frame*Stepping Limb | 34.6 | 2.3 | < 0.001 | [30.1, 39.1] |
| **Step Width** | Intercept (Walkway-fixed Frame, Inside Step) | 340.4 | 6.9 | < 0.001 | [326.8, 353.9] |
|  | Body-fixed Frame | -63.3 | 3.0 | < 0.001 | [-69.1, -57.4] |
|  | Trajectory-fixed Frame | 35.8 | 3.0 | < 0.001 | [29.9, 41.7] |
|  | Inside vs Outside | -535.6 | 3.0 | < 0.001 | [-541.5, -529.7] |
|  | Direction (CW vs CCW) | -10.2 | 1.7 | < 0.001 | [-13.6, -6.8] |
|  | Body-fixed Frame*Stepping Limb | 55.6 | 4.2 | < 0.001 | [47.3, 63.9] |
|  | Trajectory-fixed Frame*Stepping Limb | 5.6 | 4.2 | 0.184 | [-2.7, 14.0] |
| **Anteroposterior Margin of Stability** | Intercept (Walkway-fixed Frame, Inside Step) | -26.6 | 8.8 | 0.003 | [-43.9, -9.3] |
|  | Body-fixed Frame | 3.9 | 0.9 | < 0.001 | [2.0, 5.7] |
|  | Trajectory-fixed Frame | -3.5 | 0.9 | < 0.001 | [-5.3, -1.6] |
|  | Inside vs Outside | 9.2 | 0.9 | < 0.001 | [7.3, 11.0] |
|  | Direction (CW vs CCW) | -2.1 | 0.5 | < 0.001 | [-3.2, -1.0] |
|  | Body-fixed Frame*Stepping Limb | -8.9 | 1.3 | < 0.001 | [-11.5, -6.3] |
|  | Trajectory-fixed Frame*Stepping Limb | 11.7 | 1.3 | < 0.001 | [9.1, 14.4] |
| **Mediolateral Margin of Stability** | Intercept (Walkway-fixed Frame, Inside Step) | 10.6 | 3.7 | 0.004 | [3.4, 17.8] |
|  | Body-fixed Frame | -7.4 | 1.1 | < 0.001 | [-9.5, -5.2] |
|  | Trajectory-fixed Frame | 8.8 | 1.1 | < 0.001 | [6.6, 10.9] |
|  | Inside vs Outside | 127.5 | 1.1 | < 0.001 | [125.3, 129.6] |
|  | Direction (CW vs CCW) | -1.0 | 0.6 | 0.123 | [-2.2, 0.3] |
|  | Body-fixed Frame*Stepping Limb | 2.9 | 1.5 | 0.063 | [-0.2, 5.9] |
|  | Trajectory-fixed Frame*Stepping Limb | -3.3 | 1.5 | 0.033 | [-6.3, -0.3] |

**Figure 2A-D:**
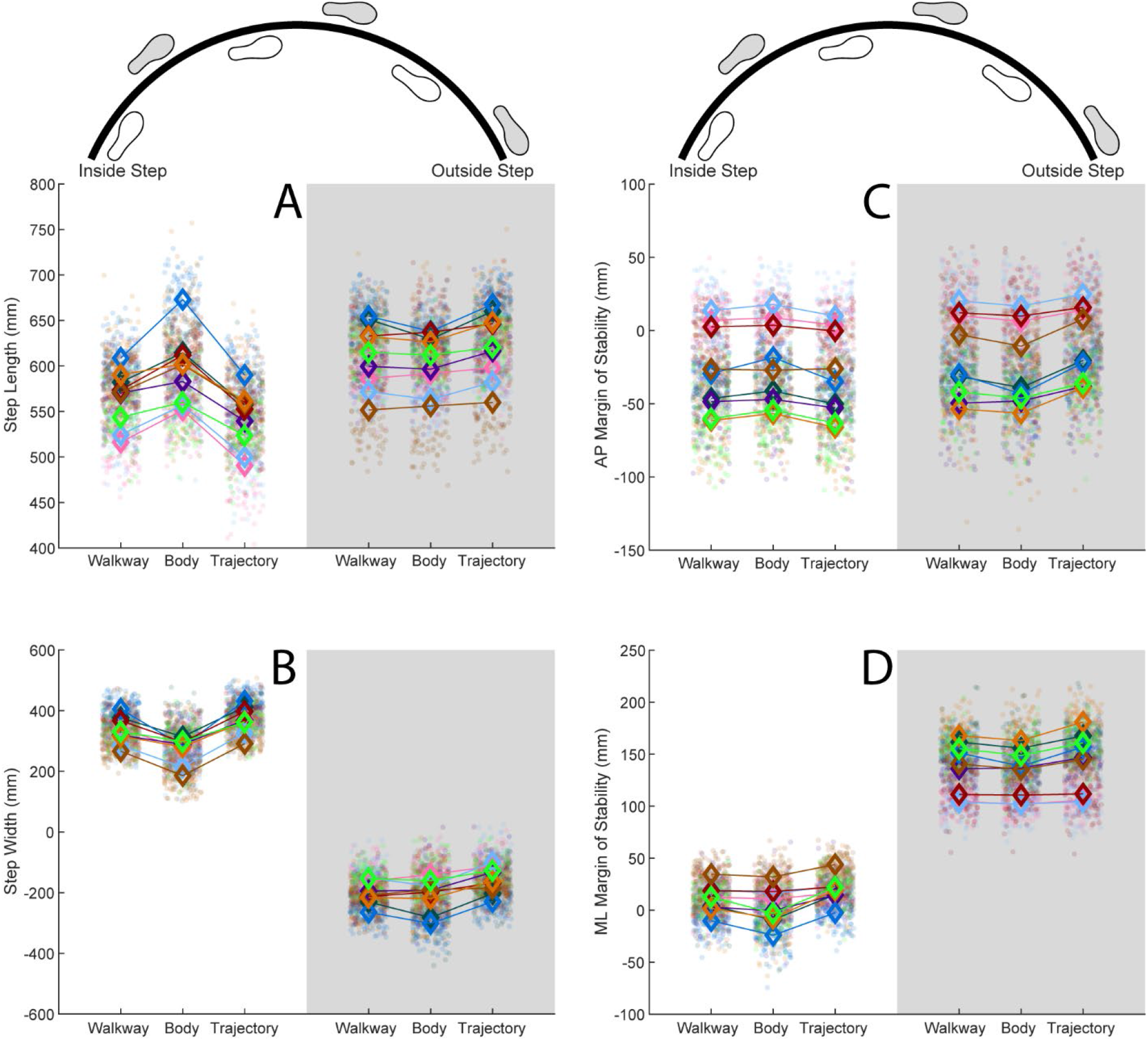
Results for each gait measure for walkway-fixed, body-fixed, and trajectory-fixed reference frames, and between inside (white) versus outside (grey) steps. Each participant’s gait measures are represented in separate colors. Each step is represented by individual scatter points. Open diamonds represent the within-participant means.

## 4. DISCUSSION

This study investigated how the choice of reference frame influences spatial measures of gait and impacts conclusions about important effects of interest (e.g., asymmetry) during a turning task. Our results indicate that different locally-based reference frames change the resulting step length, step width, AP MoS, and ML MoS. Specifically, there are large differences in all four outcome measures when comparing two common local references frames – the body-fixed frame and the trajectory-fixed frame – while the walkway-fixed frame yielded results that were between the other two definitions. For example, step width when defined in the walkway-fixed frame was smaller than when defined in the trajectory-fixed frame but greater than when defined in the body-fixed frame.

Different reference frames yielded different interpretations of asymmetry for step length and AP MoS across all three reference frames. For example, the difference in step length between inside and outside limbs was greater when using a trajectory-fixed frame compared to a walkway-fixed frame. However, the interpretation about inside vs outside steps was not different for step width in the walkway-fixed frame compared to the trajectory-fixed frame, or for ML MoS in the walkway-fixed frame compared to the body-fixed frame. While different reference frames will not universally impact every gait measure, our results indicate that any difference in reference frame will influence the interpretation of at least one outcome. Therefore, any results on measures of non-linear or turning gait should be evaluated with respect to the locally-defined reference frame used in that study. Furthermore, it is imperative that future studies investigating non-linear gait detail and justify their choice of reference frame.

When possible, we recommend using a walkway-fixed reference frame for the analysis of non-linear gait. The definition of the walkway-fixed frame is the most similar to the globally-defined reference frame that is commonly used to study linear gait (Wu and Cavanagh, 1995). Therefore, this locally-defined reference frame will likely be an intuitive choice for studies that employ a prescribed walking path. Further, the walkway-fixed frame consistently yielded results that were in-between the body-fixed and trajectory-fixed frames, indicating a desirable middle-ground. However, not all turns are attached to a specific walkway, nor are they always performed about a consistent radius (Glaister et al., 2007a; Hase and Stein, 1999), as in this study. When a walkway-fixed frame cannot be defined, a trajectory-fixed frame or a body-fixed frame can be an appropriate selection with the right justification.

Both the trajectory-fixed and body-fixed reference frames offer their own benefits and should be considered based on the outcome measures of a given study. The trajectory-fixed frame, like the walkway-fixed frame, shares some similarity to the standard globally-defined frame used in linear gait analysis; the trajectory-fixed frame is aligned with the instantaneous direction of travel, rather than the general direction of travel (Wu and Cavanagh, 1995). While similar, the instantaneous and general directions of travel differ to varying degrees throughout the gait cycle as the CoM oscillates in the ML direction during each step cycle (Tesio et al., 2010). As a result, the trajectory-fixed frame tends to oscillate between each side of the walkway-fixed frame. This oscillation within the gait cycle also makes results from the trajectory-fixed frame more dependent on the phase of the gait cycle at which they are extracted. The misalignment of the trajectory-fixed and walkway-fixed reference frames due to the ML oscillatory motion of the CoM also has important implications on any analysis of the motion of the CoM during gait, particularly regarding ML motion or ML MoS. For example, the trajectory-fixed frame is inappropriate for studies examining ML velocity of the CoM; since the trajectory-fixed frame defines the AP direction as the CoM velocity vector, the ML velocity of the CoM will necessarily be zero.

In contrast to the trajectory-fixed frame, the body-fixed frame may yield more anatomically relevant results because it is aligned with the orientation of the pelvis. Thus, a body-fixed frame may be more helpful when investigating muscle activation or joint range of motion, particularly surrounding the hip joints. However, the body-fixed frame removes a degree of freedom – axial rotation of the hip (i.e., yaw or heading angle of the pelvis) in the body-fixed transverse plane is zero at all points in time, by definition.

An important consideration for future studies of non-linear or turning gait is the calculation of margin of stability. Across a turn, individuals reorient their feet to face the new direction of travel (Bernardin et al., 2012), which can change the most anterior and most lateral points on the BoS with respect to the xCoM. These anterior-most and lateral-most points are further impacted by the definition reference frame, which establishes the AP and ML directions. Consequently, a single marker or point may not consistently represent the most lateral boundary of the BoS or effective BoS. Our study found that the most anterior boundary during every step and in each of the reference frames was the second MTP and the most lateral boundary was always the fifth MTP. But, larger turn angles with greater internal or external rotation of the foot may find different results, and we recommend considering the boundary of the BoS for each individual step, rather than using a single anatomically-fixed point, when calculating margin of stability during turning.

This study has several limitations worth noting. The primary limitation is that this study only assessed the gait of healthy young adults. It is possible that the alignment or misalignment of reference frames varies depending on different pathological gaits. However, our conclusions – that reference frames can affect results and should be explicitly defined and justified – is likely relevant to both healthy and pathological populations. Additionally, participants walked at a self-selected speed that may have altered the magnitude of the centripetal force experienced by each participant throughout the trials. It is possible that faster or slower speeds may vary the degree to which reference frames alter gait outcomes. Similarly, each trial featured the same walkway with a single consistent radius. We expect the choice of reference frame would have a greater impact on gait measures at sharper turn angles with smaller radii.

## 5. CONCLUSION

The choice of locally-defined reference frame used to measure gait during a non-linear turning task affects the results and interpretations. A walkway-fixed frame and a trajectory-fixed frame both share some similarities with the globally-defined reference frame that is commonly used in linear gait studies. But, these reference frames are not interchangeable nor synonymous. Alternatively, a body-fixed frame may provide more anatomically relevant information, but also produces different gait measures than either a walkway-fixed or a trajectory-fixed frame. Studies investigating turning gait must detail and justify their method of defining their reference frame, and results of these studies should be considered relative to their choice of reference frame.

## ACKNOWLEDGEMENTS

Special thanks to Claire Rogers for her work in pioneering the data analysis for this study, and to JunSeop Son for his assistance with data processing.

## DISCLOSURES / CONFLICTS OF INTERESTS

The authors have no conflicts of interest to declare.

## FUNDING

This work was supported by the Eunice Kennedy Shiver National Institute of Child Health and Human Development of the National Institutes of Health (award no. K12HD073945 to P.C.F.) and the University of Utah’s Undergraduate Research Opportunities Program (to TKH). Opinions, interpretations, and conclusions are those of the author and are not necessarily endorsed by the funders.

## Notes

### Competing Interest Statement

The authors have declared no competing interest.

